# A Wireless Wearable Platform for Intravenous Drug Self-Administration in Freely Behaving Rats

**DOI:** 10.64898/2026.09.08.750146

**Authors:** Eun Young Jeong, Collin D. Teague, Anisha Reimert, Sungwoo Cho, Juhyun Lee, Thorsten Althoff, Kenneth Lin, Meenakshi Nair, Tess Leong, Emily Silva, Kyle E. Parker, Catherine M. Cahill, Jordan G. McCall, Jae-Woong Jeong, Nicolas Massaly

## Abstract

Studying how drugs act on neuronal circuits requires delivering them with temporal precision while behavior proceeds undisturbed, a combination that tethered infusion systems cannot provide. We developed WEARIT (**W**ireless **E**quipment for **A**utonomous **R**at **I**nfusion **T**asks), a wearable, tetherless infusion platform that gives freely moving rats intravenous access under either remote or closed-loop operant control. The device houses a reservoir, miniaturized pump, rechargeable battery and Bluetooth circuitry in a 3D-printed enclosure worn on the back. By measuring spontaneous locomotion, amphetamine-induced hyperlocomotion and food-reinforced operant responding, we show that WEARIT leaves these behaviors unchanged. Remotely triggered fentanyl infusions yielded reliable delivery with physiological responses confirmed by pulse oximetry, and self-administration acquisition and dose–response functions were comparable to conventional tethered systems. WEARIT removes a longstanding constraint on intravenous pharmacology, opening self-administration paradigms to naturalistic and enriched environments and to concurrent imaging or optogenetic manipulation of the circuits engaged by drug reinforcement.

---

Intravenous drug delivery represents a crucial approach across pharmacology, behavioral neuroscience, and substance use disorder (SUD) research to provide rapid, precise, and reproducible control over drug exposure. Despite decades of technological advances in neuroscience, intravenous drug administration in freely behaving rodents still retains significant experimental constraints. Indeed, drugs are usually delivered either through experimenter handling, which interrupts ongoing behavior and introduces additional undesirable stress, or by using a tethered infusion system that physically connects the animal to an external benchtop drug pump. While these approaches have provided critical value in shaping our current pharmacological and circuit neuroscience knowledge, they inevitably introduce bias in behavioral outcomes. Experimenter and tethered drug infusion restrict animals’ natural movement, limit access to enriched environments and constrains the use of a variety of behavioral and neurobiological approaches. Furthermore, the use of tethers to infuse drugs imposes technical limitations in combining intravenous delivery with other modern neuroscience technologies that require optical or electrical access to the same animal.

Intravenous drug self-administration (IVSA), first established in freely moving rats by Weeks in the 1960s^1^, represents the “gold-standard” approach for investigating the behavioral, circuit, and cellular mechanisms underlying SUDs because it enables animals to voluntarily seek drug intake. However, the reliance on infusion lines has historically restricted the study of drug seeking to constrained and unenriched environments. Contexts in which drugs are sought profoundly influences drug-related behaviors. Indeed, access to alternative voluntary rewards reduces drug seeking and consumption^2–6^, while environmental enrichment decreases both opioid self-administration^7^ and reinstatement of drug seeking following abstinence^8–10^. These findings highlight the importance of studying the development and maintenance of drug seeking in more ethologically relevant environments, yet current tethered systems remain poorly suited for such investigations. To bypass the use of tethered systems many investigators have established reliable oral and nasal voluntary drug procedures including edible^11–14^, vaporized^15^ and water-based formulations^16–20^. While these approaches provide unprecedented improvements in studying untethered drug seeking, the pharmacokinetics of orally consumed and vaporized drugs are not optimal to assess accurate, time-relevant neuronal dynamics which happen concurrently with acute intravenous drug infusion, especially for drugs historically consumed intravenously by humans. Therefore, wireless intravenous delivery would remove several major technical barriers in SUD research, while enabling studies examining the impact of environmental complexity and naturalistic behaviors on drug reinforcement and the development of SUDs.

The limitations of tethered intravenous delivery extend well beyond addiction research. Recent advances in miniaturized electronics and optical technologies have transformed systems neuroscience by enabling precise interrogation of neuronal circuits during behavior^21–28^. Techniques including fiber photometry^29^, optogenetics^30^, and miniature microscopes^31,32^ now permit recordings and manipulations with unprecedented spatial and temporal resolution^26,27,33–36^, substantially advancing our understanding of circuit activity and function, and uncovering potential therapeutic targets. Despite groundbreaking progress, these approaches are rarely combined with intravenous pharmacology in freely moving animals as the use of multiple tethered connections represents substantial technical difficulties. Eliminating the intravenous tether would substantially expand the range of experimental paradigms that can easily integrate precise pharmacology with modern circuit-level neuroscience.

Lastly, remote wireless intravenous drug delivery also represents an opportunity for compounds to be delivered during uninterrupted ongoing behavior, increasing temporal precision of neuronal and behavioral responses in response to drug infusion while avoiding experimenter-induced stress or behavioral disruption. Such an approach would facilitate causal investigations of rapid drug effects during natural behavior and substantially improve the temporal alignment between pharmacological manipulations, neuronal activity, and behavioral outputs.

Here, we developed, benchmarked, and validated **W**ireless **E**quipment for **A**utonomous **R**at **I**nfusion **T**asks (WEARIT), a lightweight, wearable, Bluetooth-enabled wireless infusion platform compatible with both experimenter-triggered and operant self-administration intravenous drug delivery in freely behaving rats. Using physiological and behavioral validation experiments, we demonstrate that WEARIT does not alter locomotion or operant reward seeking, reliably delivers intravenous bolus injections, and supports chronic opioid self-administration with drug intake levels comparable to conventional tethered IVSA systems. By enabling intravenous drug delivery in unrestricted animals without interrupting ongoing behavior, WEARIT provides a versatile platform that expands the experimental possibilities for pharmacology, systems neuroscience, and addiction research.

## RESULTS

### WEARIT enables wireless intravenous drug delivery in freely behaving rats

WEARIT is a wireless intravenous drug infusion platform designed to be worn on the dorsum as a “backpack” and secured with an adjustable wearable Velcro strap (**Fig. 1a-c**). A lightweight 3D-printed enclosure (**Supplementary Fig. 1**) houses the fluidic and electronic components required for intravenous infusion while permitting unrestricted locomotion (**Fig. 1b,c**). The assembled system measures 26 × 26 × 51 mm³ and weighs 16.8 g, corresponding to approximately 2.8-5.6% of the body weight of a typical adult rat (300-600 g), within the range typically considered tolerable for dorsal backpack-mounted devices^37^. The micropump draws drug solution from the onboard reservoir (**Fig. 1c**) and delivers it through silicone tubing and a Luer-lock connector to an indwelling jugular catheter (**Fig. 1b and Supplementary Fig. 2**), establishing a compact fluidic pathway from the wearable reservoir to the intravenous access port. The hand-made refillable drug reservoir (**Supplementary Fig. 3**), holding up to 5 mL of solution, adds minimal (∼5 grams) weight to the overall device when filled up. While the dimensions of the reservoir can be adjusted for specific experimental needs, the added weight from drug solution should be considered as potential source of locomotor burden.

**Fig. 1.**
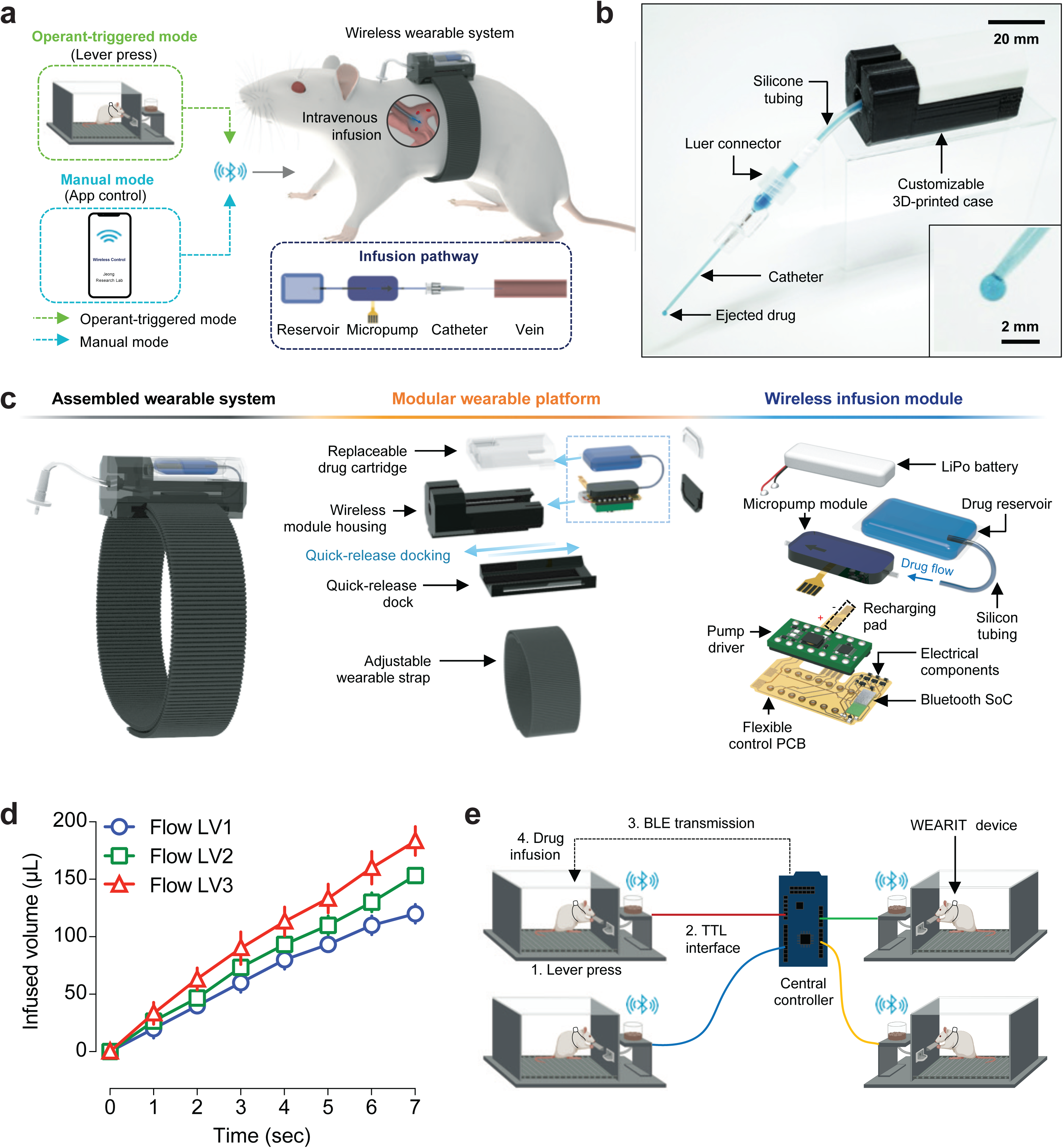
Concept and design of WEARIT for wireless intravenous drug delivery in freely behaving rats. **a**, Schematic representation of the WEARIT platform highlighting two wireless operating modes: an operant-triggered mode, in which a lever press initiates infusion, and a manual mode, in which infusion is triggered remotely through a Bluetooth-enabled interface. The lower-right inset shows the drug delivery path from the onboard reservoir, through the micropump and catheter, to the jugular vein. **b**, Photograph of the WEARIT fluidic interface, including the customizable 3D-printed enclosure, silicone tubing, Luer-lock connector and catheter. Inset shows a magnified view of drug ejection from the catheter tip. **c**, Modular design of WEARIT. <u>Left</u>: assembled wearable system mounted on an adjustable strap. <u>Center</u>: exploded view of the modular wearable platform, including a quick-release dock, a wireless module housing and a replaceable drug cartridge. <u>Right</u>: exploded view of the removable wireless infusion module, comprising the LiPo battery, micropump module, drug reservoir and a flexible control PCB that integrates the Bluetooth System-on-Chip (SoC), pump driver and recharging pad. **d**, Infused volume as a function of time for three software-selectable flow-rate settings (Flow LV1: 1.05 mL min^-1^, LV2: 1.26 mL min^-1^, LV3: 1.55 mL min^-1^), demonstrating programmable control of drug delivery (mean ± s.d.; *n* = 3 devices per setting). **e**, Schematic representation of the operant-triggered wireless control architecture. A lever press generates a transistor–transistor logic (TTL) signal ①, which is relayed through a TTL interface to a central controller ②, converted into a Bluetooth Low Energy (BLE) command and transmitted wirelessly to the designated WEARIT device ③, thereby triggering infusion in the corresponding animal ④.

Importantly, WEARIT adopts a modular architecture that permits rapid attachment, removal, and replacement of the wireless infusion module without anesthetizing the animal (**Fig. 1c**). The wireless module housing, assembled with the replaceable drug reservoir, interfaces with a quick-release dock through a U-shaped guide rail, enabling module exchange with minimal procedural burden. The drug reservoir is fabricated from heat-sealed polyethylene material with silicone tubing integrated at the fluidic interface, yielding a low-cost, flexible, adjustable and replaceable cartridge (**Supplementary Fig. 3**). A recharging pad integrated onto the flexible printed circuit board enables battery charging through a custom-made seven-channel charging board without disassembling the enclosure (**Supplementary Fig. 2 and 4**), supporting repeated use across multiday behavioral experiments.

The wireless infusion module provides programmable control of infusion rate and volume through software-controlled selection of discrete pump-driving levels (**Supplementary Fig. 5**). Infused volume increased linearly with infusion duration across all tested flow-rate settings, confirming a predictable control of delivered volume by pump activation time (**Fig. 1d**). A central wireless controller board (nRF52840-PCA10056, Nordic Semiconductor) allows for coordination of multiple WEARIT devices in parallel for high-throughput operant self-administration experiments (**Fig. 1e**). Transistor–transistor logic (TTL) signals generated from active lever presses from individual operant chambers are routed to a single wireless central controller board, which dispatches device-specific Bluetooth Low Energy (BLE) commands to trigger infusion in the designated animal (**Fig. 1e**). This architecture enables independent and selective wireless control of intravenous delivery across multiple animals without tethered infusion lines or a dedicated external pump for each chamber.

### WEARIT provides precise fluidic control and scalable wireless triggering

Reliable wireless communication and precise drug delivery are essential prerequisites for untethered IVSA experiments. WEARIT uses a piezoelectrically actuated micropump with passive check valves that enforce unidirectional fluid flow during cyclic membrane deformation, enabling drug transport from the onboard reservoir to the implanted catheter (**Fig. 2a**). Stepwise increases in the pump-driver voltage produced three discrete flow-rate levels of 1.05, 1.26, and 1.55 mL min^-1^, respectively, providing a programmable control of infusion rate (**Fig. 2b**). Dosing accuracy was assessed by comparing programmed target volumes with experimentally measured outputs. Delivered volumes deviated from targets by less than 0.3 μL across all conditions, yielding mean measured values of 24.93, 50.01, 75.20, and 99.79 μL, respectively (*n =* 10 per condition) (**Fig. 2c**).

**Fig. 2.**
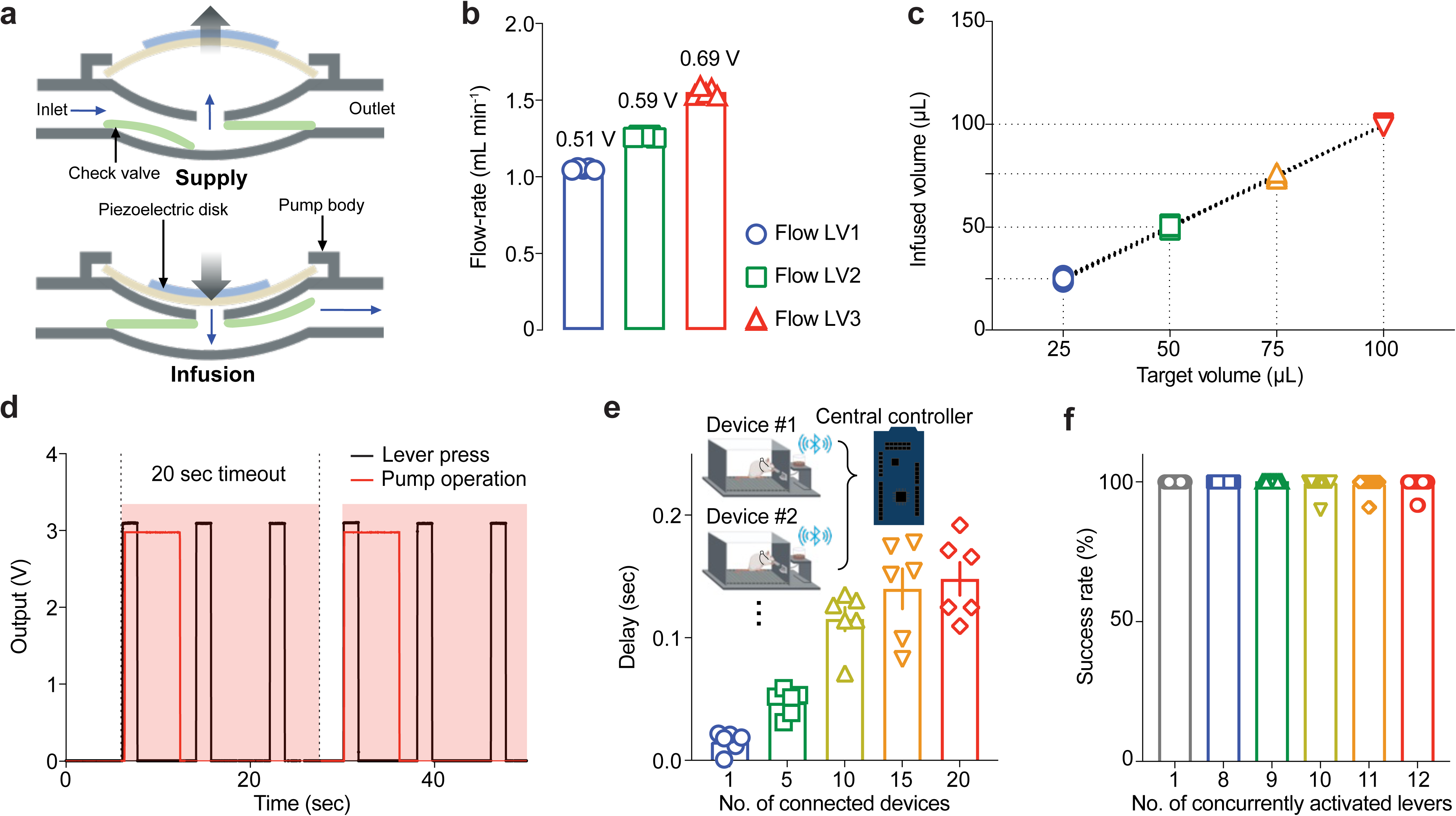
Fluidic performance and wireless communication characteristics of WEARIT. **a**, Operating principle of the piezoelectrically actuated micropump. Cyclic deformation of the piezoelectric disk drives fluid from the inlet to the outlet, with passive check valves maintaining unidirectional flow across the supply and infusion phases. **b**, Flow-rate of WEARIT at three software-selectable pump-driving levels (Flow LV1, 1.05 mL min^-1^; Flow LV2, 1.26 mL min^-1^; Flow LV3, 1.55 mL min^-1^), demonstrating programmable control of infusion rate (mean ± s.e.m.; *n* = 5 devices). **c**, Quantification of the delivered infusion volume as a function of expected target volume. This shows accuracy between programmed and delivered volumes (mean ± s.e.m.; *n* = 10 trials). **d**, Timing trace of an operant-triggered infusion sequence. Lever-press events (black) triggered micropump activation (red). A 20-s lockout period was implemented after each activation to prevent unintended re-dosing. **e**, Wireless communication delay as a function of the number of WEARIT devices connected to a single central controller. Delay was defined as the interval between a lever-press and pump activation in the triggered device (mean ± s.e.m.; *n* = 6 trials). **f**, Pump activation success rate as a function of the number of concurrently activated levers, with 20 WEARIT devices connected to a single central controller condition (mean ± s.e.m.; *n* = 20 trials per condition).

We next evaluated whether this performance was maintained under repeated-use conditions, mirroring IVSA experiments. Across 50 consecutive trials, delivery remained stable at each programmed volume, confirming infusion-to-infusion reproducibility (**Supplementary Fig. 6a**). Using the onboard 5 mL drug reservoir with a 100 μL target volume per infusion, delivery accuracy was maintained within 2.67% of the target through 48 consecutive infusions, after which accuracy declined as the drug supply became scarce (**Supplementary Fig. 6b**). The total device power reached only 33.19 mW at the highest flow-rate setting (**Supplementary Fig. 7a,b)**, and a fully charged 120-mAh battery sustained device operation for 13 days under a simulated IVSA schedule consisting of daily 1-h sessions with twelve 50 μL infusions delivered at Flow LV2 at 5-min intervals (**Supplementary Fig. 7c,d**). Together, these benchtop measurements demonstrate accurate, reproducible and energy-efficient infusion control suitable for longitudinal behavioral experiments.

To support untethered operant experiments, WEARIT translates lever-press events into wireless pump activation commands in real time. After each infusion, a 20-s lockout period was established to suppress additional lever-press-triggered infusions and prevent unintended re-dosing (**Fig. 2d**). As each WEARIT device carries a unique identifier, the central controller board can selectively activate a designated device among multiple simultaneous connections. Communication delay increased modestly as more devices connected to the central controller, but the interval between lever-press detection and pump activation remained below 0.2 s across all tested conditions, including simultaneous BLE connections to 20 WEARIT devices from a single central controller (**Fig. 2e**). Scaling to a larger number of connected devices, according to experimental needs, can be supported by adding more central controllers^25,28,38^. Under conditions emulating concurrent lever presses across multiple operant chambers, pump-activation success rates remained above 99%, reaching 99.17% even when 12 levers were triggered simultaneously (**Fig. 2f**). Wireless triggering remained reliable over distances of up to 50 m, with no transmission failures observed under line-of-sight conditions (**Supplementary Fig. 8**). Together, these results demonstrate that WEARIT supports low-latency, device-specific wireless triggering across scalable arrays of behavioral chambers.

### WEARIT does not disrupt spontaneous locomotion

To determine whether WEARIT devices could be implemented during freely moving behavioral tasks, we measured spontaneous locomotor behavior in an open field test using both male and female rats. Animals were first habituated to carrying the WEARIT device for 20 minutes per day for five consecutive days. During these habituation procedures, WEARIT was attached to a standard vascular access harness on the rat’s back before placing the animal in a dimly lit open field (**Fig. 3a**). On the sixth day, animals were placed back in the open field either untethered or carrying the WEARIT device attached to their harness and their horizontal locomotion was monitored for 20 minutes using video tracking software (**Fig. 3a**). Across the 20-minute session, both male and female rats equipped with WEARIT exhibited locomotor activity comparable to that observed in untethered harness-only controls (**Fig. 3b,c**). While the cumulative distance traveled across time did not differ significantly (males: *P* = 0.3429, females: *P* = 0.40), a slight and non-significant decrease in overall activity could be observed. While underpowered to test selective sex difference, locomotor activity was not impacted in either sex regardless of whether the rats carried the WEARIT device on their back (**Fig. 3d,e**).

**Fig. 3.**
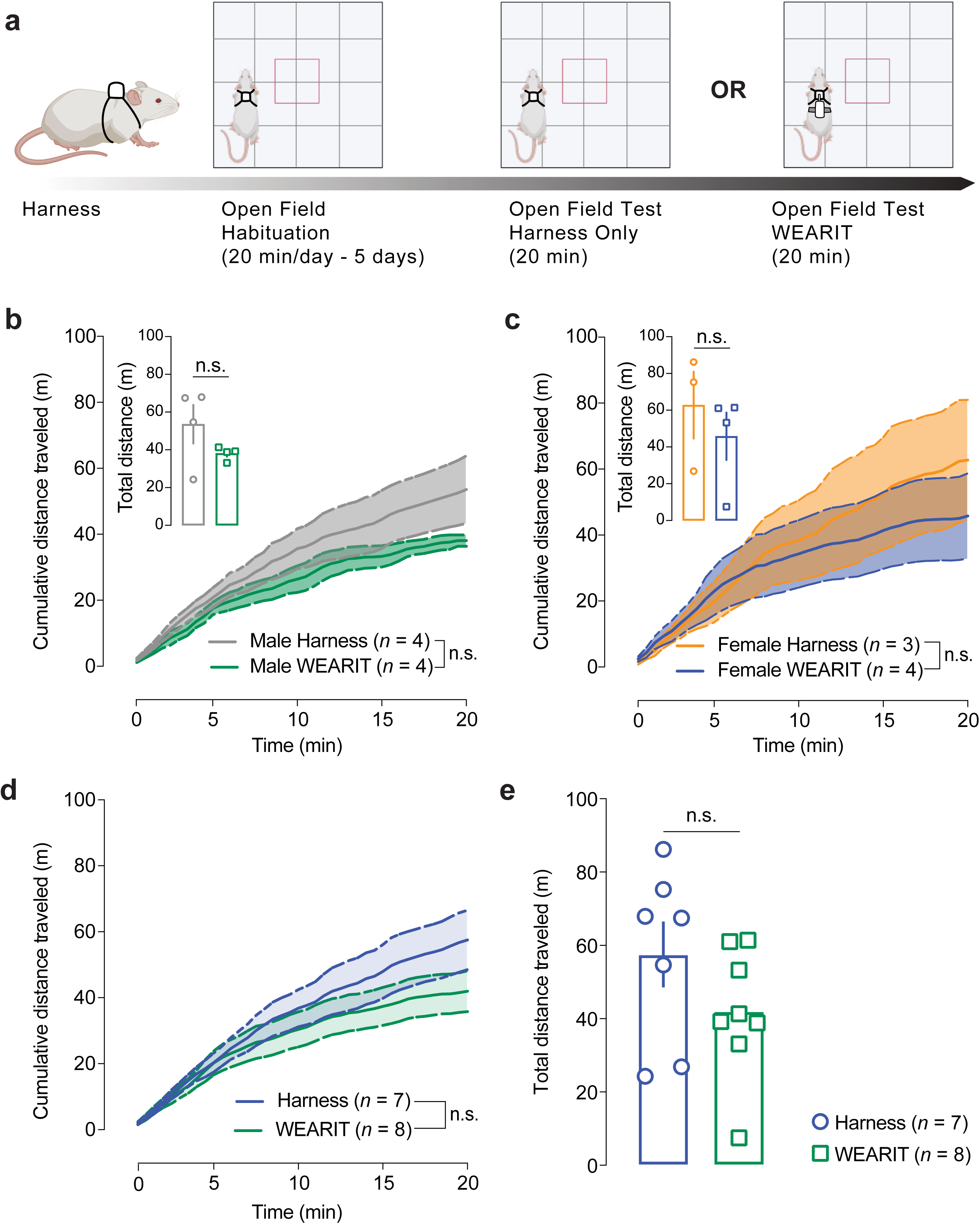
WEARIT does not impair locomotor activity. **a**, Schematic representation of the behavioral methodology. All animals were habituated to carry the WEARIT on the back for 20 min per day for 5 days. The following day rats were either equipped with the WEARIT or left unequipped (Harness group) before being placed in the open field arena. Locomotor activity was measured for 20 min. **b–c**, Cumulative distance traveled across the 20 min session does not differ between WEARIT and control (Harness) groups for both male (**b**, Harness *n* = 4, WEARIT *n* = 4) and female (**c**, Harness *n* = 3, WEARIT *n* = 4) rats. Insets demonstrate that the overall distance traveled during the 20 minutes session does not significantly differ between control (Harness) and WEARIT groups. **d**, Cumulative distance traveled across the 20 min session does not differ between WEARIT and control (Harness) groups when male and female rats are combined (Harness *n* = 7, WEARIT *n* = 8). **e**, The overall distance traveled during the 20-minute session (mean ± s.e.m.) does not significantly differ between control (Harness, *n* = 7) and WEARIT (*n* = 8) groups. All data shown as mean ± s.e.m. and n.s. *P* > 0.05.

To further investigate a potential impact of WEARIT on locomotor activity, we assessed whether WEARIT would impact psychostimulant-induced hyperlocomotion. In this set of experiment, rats were habituated to intraperitoneal injections (sterile saline, 1 mL kg^-1^) and directly placed in a dimmed light open field for 30 minutes for three consecutive days (**Fig. 4a**). On the fourth day, rats received a single acute infusion of amphetamine (2 mg kg^-1^, i.p.) and were placed in the open field, either untethered (control) or carrying the WEARIT device. A significant impact of the device on locomotor activity could be observed during habituation sessions for both males and females (**Fig. 4b,c**; two-way ANOVA for repeated measures: <u>Males</u>: WEARIT vs Control, F_(1,10)_ = 6.128, *P* = 0.0328, <u>Females</u>: WEARIT vs Control, F_(1,10)_ = 72.29, *P* < 0.001). While the WEARIT devices only produced a non-significant decrease in male rats locomotion on the third day of habituation (distance traveled: 80.89 m ± 4.612 vs 54.84 m ± 9.108 in control vs WEARIT respectively, *P* = 0.1140, *n* = 6, Sidak post hoc test), this difference was striking in female rats (distance traveled: 44.52 m ± 5.708 vs 129.9 m ± 9.826 in control vs WEARIT respectively : *P* < 0.0001, *n* = 6 Sidak post hoc test) (**Fig. 4c**). This result demonstrates the necessity for extended habituation to the devices to buffer difference in ambulatory behavior. Lastly, the WEARIT does not significantly decrease amphetamine-induced hyperlocomotion in males (*P* = 0.1996, *n* = 6, two-tailed paired t-test) or in females (*P* = 0.1697, *n* = 6, two-tailed paired t-test) (**Fig. 4d,e**). Further, a single acute infusion of amphetamine significantly increases locomotor compared to saline (data from the last habituation session) in both control (*P* = 0.0012, *n* = 12 Sidak post hoc test) and WEARIT (*P* = 0.0004, *n* = 12 Sidak post hoc test) groups, confirming that WEARIT does not impair our ability to measure psychostimulant-induced hyperlocomotion.

**Fig. 4.**
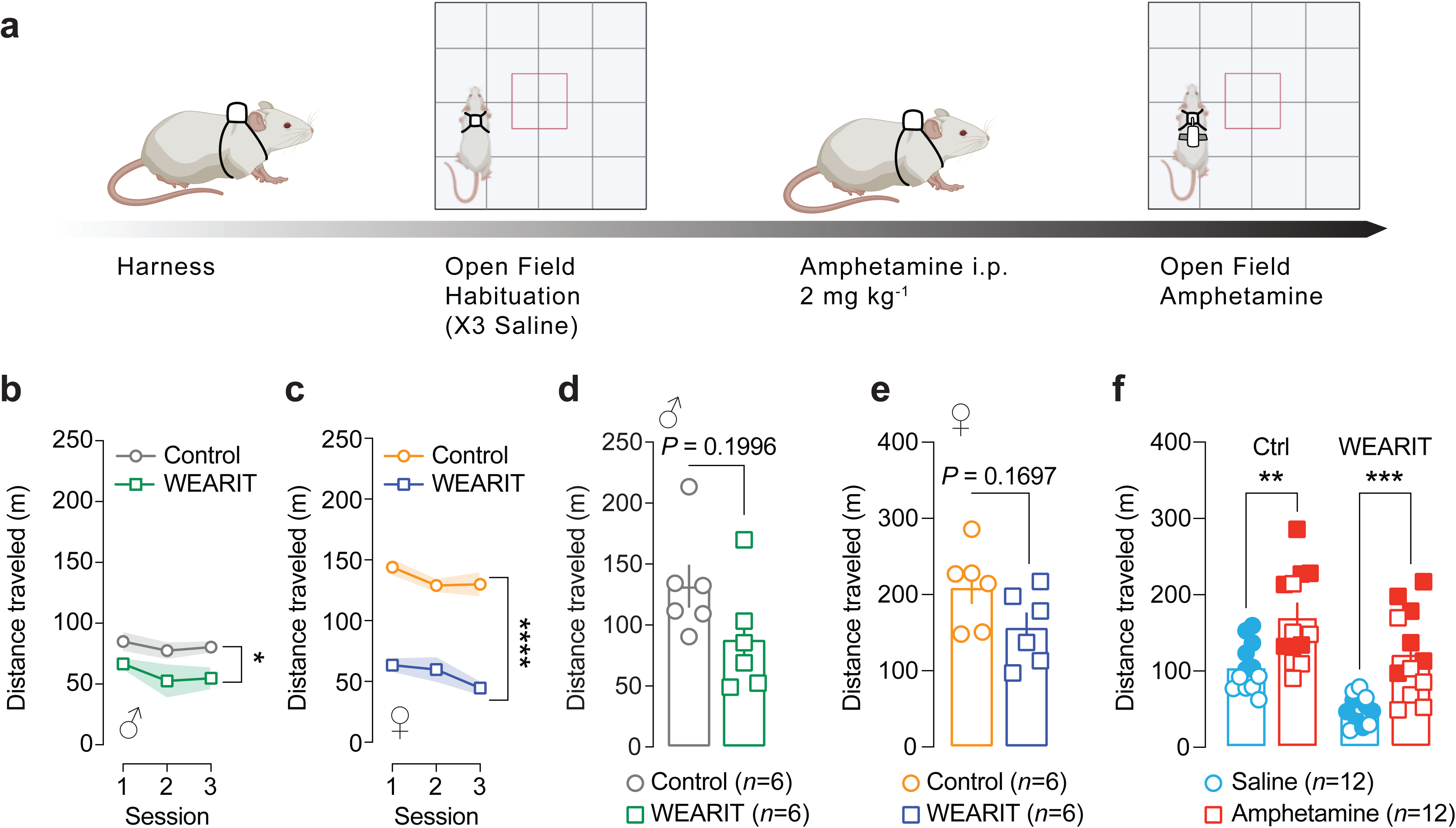
WEARIT does not impair stimulant-induced increases in locomotor activity. **a**, Schematic representation of the behavioral methodology. All animals were habituated to carry the WEARIT on the back for 20 minutes per day for three days following a Saline intraperitoneal injection. The following day rats were injected with amphetamine (2 mg kg^-1^, i.p.) and either equipped with the WEARIT or left unequipped (Harness group) before being placed in the open field arena directly after drug infusion. Locomotor activity was measured for 20 minutes. **b–c**, WEARIT does significantly reduce locomotor activity during habituation sessions in both male (**b**) and female (**c**) rats (*n* = 6 per group). **d–e**, The overall total distance traveled during the 20-minute locomotor session after amphetamine infusion does not differ between harness and WEARIT group for both male (**d**, *n* = 6) and female (**e**, *n* = 6) rats. **f**, Amphetamine infusion significantly increases locomotor activity in both control and WEARIT groups compared to the last habituation day after saline injection (**f**, n=12 per group, full and empty symbols represent females and males, respectively). All data shown as mean ± s.e.m. * *P* < 0.05; ** *P* < 0.01; *** *P* < 0.001; **** *P* < 0.0001.

Altogether, these findings indicate that, with optimal habituation sessions, WEARIT does not impair spontaneous or psychostimulant-induced locomotor activity. These data demonstrate that the WEARIT device does not introduce substantial disturbances to ongoing and drug-evoked behaviors, allowing for measurements of drug-selective neuronal and behavioral responses to real-time remote drug infusion.

### WEARIT does not impair operant responding or reward consumption

In addition to ambulatory behavior, rats’ ability to interact with their environment, collect and consume rewards such as food pellets remain critical for assessing drug and reward seeking and reinforcement. Here, we assessed whether WEARIT devices impair rats’ interactions with operant levers and food consumption in commercially available operant boxes (**Fig. 5a**). First, rats were trained to acquire sucrose pellets for 5 days under a Fixed-Ratio 1 schedule of reinforcement (FR1) during which an active lever press delivered a 45 mg chocolate flavored food pellet (**Fig. 5b**). No difference was observed in learning between the groups (two-way ANOVA for repeated measures. WEARIT vs Control: F_(1,16)_ = 0.01877, *P* = 0.8927 for pellets obtained, and F_(1,16)_ = 0.1724, *P* = 0.6835, for active lever presses). Inactive lever presses, used as control for learning, remained low and similar across groups (two-way ANOVA for repeated measures. WEARIT vs Control: F_(1,16)_ = 0.03824, *P* = 0.8474), indicating discrimination between active and inactive lever interactions. After each operant conditioning session, rats were habituated to carrying the WEARIT device on their back 15 minutes per day. A day after operant food seeking was acquired (no more than 10% variation in pellets obtained across three consecutive days), the potential impact of WEARIT disturbances on food seeking and consumption was assessed using an additional FR1 session.

**Fig. 5.**
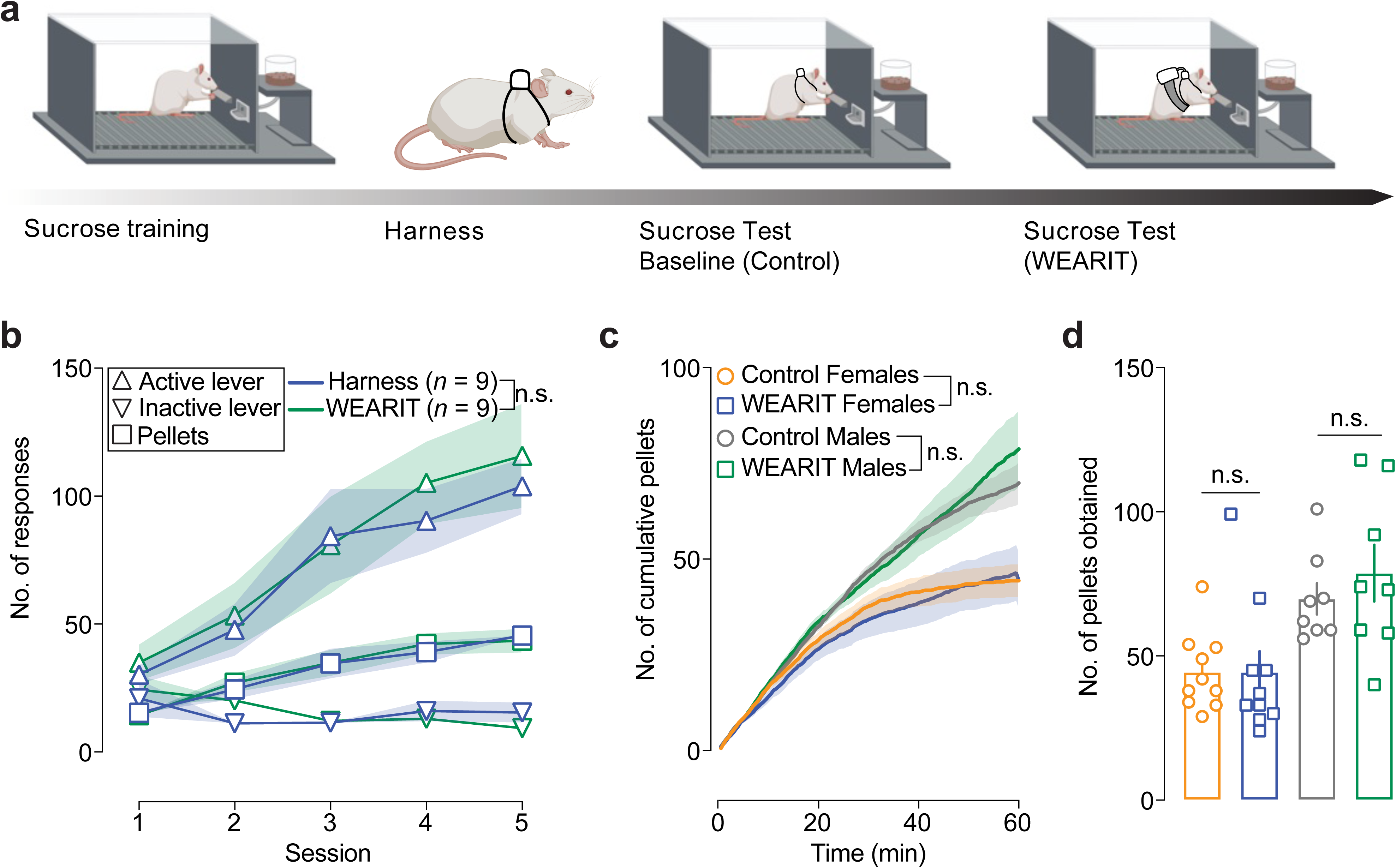
WEARIT does not impair operant reward seeking and consumption. **a**, Schematic representation of the behavioral methodology. Briefly, all animals were exposed to 5 days of sucrose self-administration training under a Fixed Ratio 1 (FR1) schedule of reinforcement. After harness placement and habituation session for the WEARIT device, rats were exposed to a control one-hour operant sucrose self-administration session during which they would not carry the WEARIT device. The day after, the same animals were re-exposed to a one-hour sucrose self-administration session after the WEARIT device was placed on their back. Order of control and WEARIT tests were counterbalanced so half of the animals ran the WEARIT session on the first day and half of the animals ran the WEARIT session on the second day. **b**, All animals learn sucrose self-administration procedure across five days of training. **c**, Male and female rats display similar rates of pellets acquisition across time during both control and WEARIT sessions. **d**, The total number of pellets obtained during the one-hour sucrose self-administration session does not differ between control and WEARIT groups for both male and female rats. All data shown as mean ± s.e.m. and n.s. *P* > 0.05.

During this test session, animals equipped with WEARIT exhibited levels of active lever responding comparable to untethered control animals (two-way ANOVA for repeated measures. WEARIT vs Control: <u>Males</u>: F_(1,14)_ = 0.05694, *P* = 0.8149, and <u>Females</u>: F_(1,18)_ = 0.1217, *P* = 0.7313 across the session (**Fig. 5c**). In addition, the number of pellets obtained, and the cumulative time course of pellet retrieval were indistinguishable between groups (<u>Females</u>: two-tailed Wilcoxon matched-pairs signed rank test: *P* = 0.4317; and <u>Males</u>: two tailed t-test for paired values: *P* = 0.5156) (**Fig. 5c,d**). These results demonstrate that the WEARIT infusion system does not impair the animals’ ability to interact with operant devices or retrieve and consume food rewards. Together, these data establish that the WEARIT platform is compatible with behavioral paradigms requiring precise operant responding, enabling investigators to further assess the precise temporal effects of intravenous drug infusion during ongoing reward seeking in unrestricted animals.

### WEARIT provides reliable intravenous drug infusion

Having established that the wearable system does not disrupt baseline behavior, we next tested whether remotely-infused drugs delivered through the WEARIT platform reliably enter the bloodstream and produce expected physiological effects. To reach this goal, rats were first permanently implanted with an intravenous catheter in their jugular vein connected to a wearable harness (VAHR1/22, Instech). After five days of habituation to the WEARIT device, rats received experimenter-triggered wireless fentanyl infusions (50 μg kg⁻¹), or sterile saline as control. Respiratory parameters were monitored using a non-invasive tethered pulse oximetry system (Starr Life Sciences) to assess physiological consequences of fentanyl infusion (**Fig. 6a**). Fentanyl delivery produced a rapid and robust respiratory depression compared to sterile saline injections (Two-way ANOVA for repeated measures: F_(1, 10)_ = 31.20; *P* = 0.0002), consistent with the well-described respiratory depressant effects of mu opioid receptor agonists (**Fig. 6b**). Arterial blood oxygen saturation levels decreased following fentanyl infusion, confirming reliable intravenous drug delivery.

**Fig. 6.**
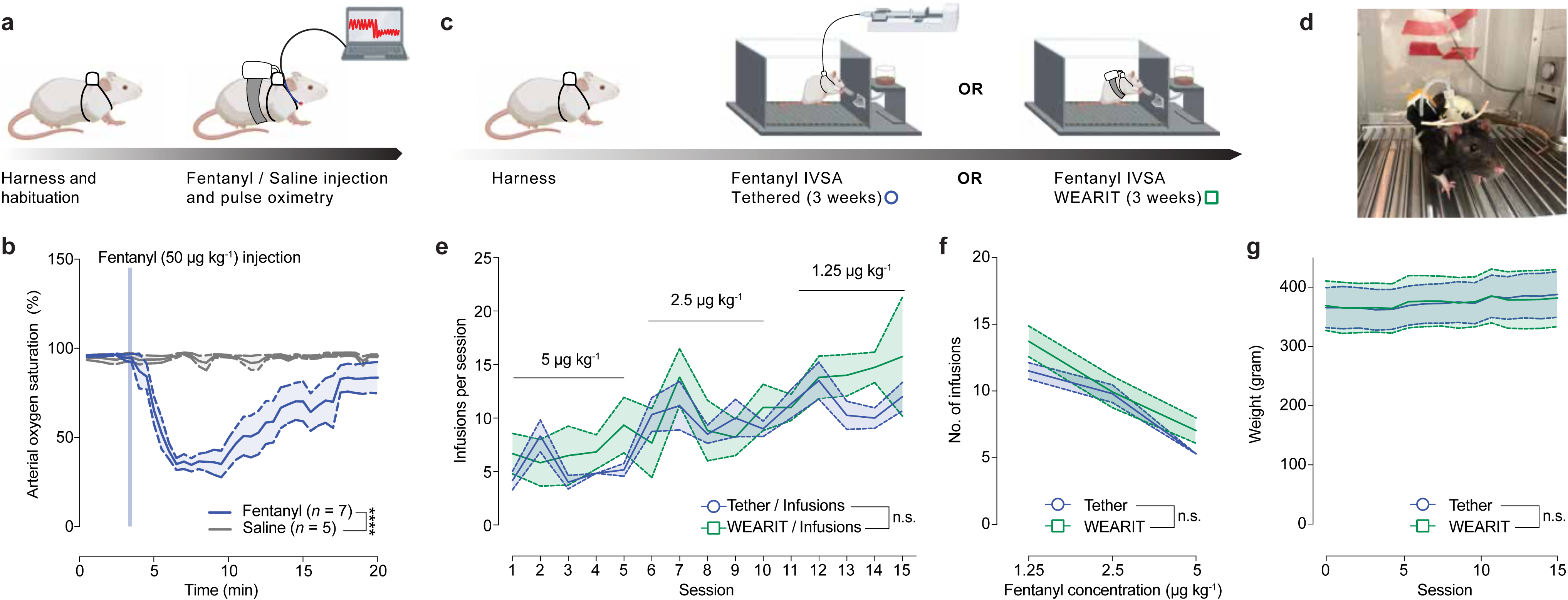
WEARIT allows for intravenous drug delivery and operant intravenous self-administration procedures. **a**, Schematic representation of the behavioral methodology. After catheter implantation procedure animals were habituated to the WEARIT and non-invasive pulse oximeter collar devices for five days. The following day, animals received an experimenter-triggered intravenous fentanyl infusion (50 µg kg^-1^, *n* = 7), or saline as control (*n* = 5). Arterial oxygen saturation was monitored as a readout for the respiratory depressant effects of fentanyl. **b**, Rats injected with fentanyl demonstrate a rapid and significant decrease in oxygen saturation following drug infusion, compared to saline injected rats (*** *P* < 0.001). The effect persists for up to 15 minutes before returning to baseline levels. **c**, Schematic representation of the behavioral methodology. After catheter implantation procedure animals were habituated to the WEARIT for 5 days. They were exposed to 5 sessions per week of fentanyl self-administration under fixed ratio 1 (FR1) schedule of reinforcement using either a classic tethered or WEARIT approach. Each week the concentration of the fentanyl delivered intravenously was decreased by half to assess whether animals would increase the number of infusions obtained per session to counterbalance lower drug concentrations. **d**, Representative photo of a rat carrying the WEARIT device on its back. **e**, Number of infusions / lever presses per session did not differ between tethered (*n* = 6) or WEARIT (*n* = 6) groups across the three weeks of intravenous self-administration. **f**, Similarly, the mean number of infusions per session obtained for each dose assessed during our dose-response experiment did not differ between tethered and WEARIT groups. **g**, The overall rat weight (in grams) for each group did not differ across the 3 weeks experiment. All data shown as mean ± s.e.m. *** *P* < 0.001, and n.s. *P* > 0.05.

### WEARIT enables fentanyl self-administration comparable to tethered IVSA

Finally, we tested whether operant fentanyl self-administration could be achieved with WEARIT. To do so, we benchmarked fentanyl seeking and consumption in WEARIT to conventional tethered IVSA paradigms. A week after receiving intravenous catheter implantation and habituation to WEARIT devices, rats were placed in operant boxes and trained to self-administer fentanyl for one hour daily using either a standard tethered infusion system or the WEARIT wireless platform (**Fig. 6c**). Each WEARIT was individually controlled using a wireless central controller board. Across all training sessions, rats equipped with WEARIT displayed levels of active lever responding and infusion intake comparable to tethered controls (Two-way ANOVA for repeated measures – mixed effects analysis: WEARIT vs Tether: F_(1,10)_ = 1.107, Interaction WEARIT vs Tether x time: F_(3.574, 32.68)_ = 0.8694; **Fig. 6d**). Further, after a week of operant fentanyl seeking and consumption, the concentration of fentanyl solution was decreased to assess whether rats would increase the number of fentanyl doses obtained per session to counter the lower concentration of the solution (**Fig. 6e,f**). Both groups increase their intake similarly after fentanyl concentration decreased twice, demonstrating that dose–response testing across multiple fentanyl concentrations produced similar intake patterns across delivery methods (Two-way ANOVA – mixed effects analysis: WEARIT vs Tether: F_(1,58)_ = 3.143, Interaction WEARIT vs Tether x time: F_(1.698, 88.31)_ = 0.8494; **Fig. 6e**). As rats kept a steady weight across the whole experiment, the observed increase in number of drug infusions per session could not be accounted to a higher body weight (**Fig. 6g**).

Together these results demonstrate that the WEARIT platform supports IVSA behavior and closely matches drug seeking and intake observed using conventional tethered IVSA procedures enabling further work in freely moving rats in complex environments.

## DISCUSSION

In the present study we developed WEARIT, a wireless wearable platform for intravenous drug delivery. WEARIT enables experimenter-triggered or self-administered drug infusions in freely moving rats without tethering. Using a combination of physiological recordings and behavioral validation experiments, we demonstrate that WEARIT does not alter spontaneous locomotion or operant responding in freely moving rats, reliably delivers drug intravenously, and allows reliable untethered IVSA. By removing the constraints inherent to tethered intravenous infusions, WEARIT enables temporally precise intravenous pharmacology in undisturbed, freely behaving animals, establishing a technological platform broadly applicable across neuroscience, pharmacology, and translational drug discovery.

One immediate advantage of the platform is the ability to combine intravenous drug delivery with modern systems neuroscience approaches. As integrating modern tethered approaches and intravenous pharmacology has remained technically challenging, investigators often resort to either running IVSA in movement-restricted or head-fixed setups^39,40^, or use alternative routes of administrations for voluntary seeking and consumption^11–20^. By eliminating the need for a tether to infuse drugs, WEARIT enables the integration of IVSA and other tethered approaches in freely moving animals, as shown with our non-invasive pulse oximeter collars (**Fig. 6a,b**). We anticipate that this compatibility will empower the field to further investigate the neuronal mechanisms underlying acute intravenous drug seeking, consumption, and relapse-related behaviors using a route of administration relevant to intravenously consumed substances.

Beyond compatibility with existing recording technologies, WEARIT fundamentally improves the temporal precision with which intravenous pharmacology can be studied. Experimenter-administered drugs often require approaching or handling the animal immediately before drug delivery, disrupting ongoing behavior and introducing stress that confounds both behavioral and neuronal responses to the drug. In contrast, WEARIT enables investigators to acquire temporally precise and relevant information on discrete neuronal populations selectively engaged by intravenously delivered compounds during ongoing behavior. Such temporal precision is likely to be valuable for experiments combining intravenous pharmacology with functional MRI, microPET imaging, calcium imaging and other approaches aimed at understanding the rapid actions of drugs on neuronal circuits and networks.

The WEARIT platform also substantially expands the behavioral contexts in which intravenous drugs can be studied. Conventional IVSA paradigms have historically been confined to relatively simple operant chambers ^3,9,41^. Growing evidence demonstrates that environmental complexity, access to alternative rewards, and social interactions profoundly impacts drug seeking, consumption, and relapse^2,8,9,42,43^. WEARIT now makes it feasible to develop intravenous self-administration paradigms in enriched and naturalistic environments while simultaneously monitoring or manipulating neuronal activity. Such experiments should provide a more comprehensive understanding of how environmental context shapes drug reinforcement and may improve the translational relevance of preclinical models of substance use disorders.

The engineering design of WEARIT necessarily reflects a series of tradeoffs between functionality, device mass, battery life, and drug availability within a session. We deliberately optimized the platform to remain lightweight, with the complete wearable system remaining well below the commonly accepted 10% threshold for backpack-mounted devices. Within these constraints, WEARIT integrates a ∼5 mL refillable drug reservoir, a programmable infusion pump, a rechargeable battery, and a Bluetooth communication module while maintaining normal spontaneous locomotion and operant behavior in rats.

With these constraints, the current WEARIT platform was specifically designed for rats and is not directly translatable as a wearable system for mice due to their smaller weight and size. Nevertheless, the wearability of our platform is only one component of the overall technology. Using a combination of the programmable pump and a centralized wireless controller can readily be adapted as a compact and inexpensive (∼$250) standalone infusion system for tethered mouse experiments (**Supplementary Table 1**). Such an implementation could provide an attractive alternative to conventional syringe-pump systems and would be readily compatible with commercially available operant platforms as well as the open-source FED3 system^44^, while substantially reducing experimental cost and infrastructure requirements.

Our housing module was also designed with experimental flexibility in mind. The quick-release docking mechanism allows investigators to exchange reservoirs, batteries, or complete infusion modules within seconds without anesthetizing the animal. Although the relatively limited capacity of the onboard reservoir currently restricts the duration of continuous self-administration sessions, our design enables rapid replacement of prefilled reservoirs during long-access paradigms while minimizing interruption of the experiment.

An additional limitation of the current platform is that it was designed primarily for individually housed animals. Because the external device remains accessible to conspecifics, social housing would likely produce mechanical damage from cage mates. This limitation is particularly important as social interactions can significantly decrease drug seeking, consumption, and relapse vulnerability^2,8,9,45^. Future generations of WEARIT could incorporate rodent-resistant protective jackets or integrated protective housing that permit long-term group housing without compromising device integrity. Coupling such systems with RFID-based tracking technologies and automated IoT-enabled systems in large, enriched environments^25,28,38,46^ would represent an important next step toward establishing intravenous self-administration paradigms in socially housed animals and allow for assessing how social hierarchy and environmental complexity could promote vulnerability or resilience to SUDs.

Importantly, the utility of WEARIT is not restricted to substance use disorders and neuroscience research. Many experimental and therapeutic compounds are administered intravenously during preclinical studies to characterize pharmacokinetics, efficacy, and behavioral effects. WEARIT offers new opportunities for evaluating candidate analgesics, anesthetics, psychiatric therapeutics, cardiovascular drugs, and/or immunomodulators and investigating the behavioral consequences and rapid central or peripheral actions of these pharmacological interventions.

Lastly, WEARIT upgrades intravenous drug delivery into a programmable variable. Because our drug infusion system is software-defined, Bluetooth-enabled, controls individual devices independently, and is triggered with sub-second latency, WEARIT could be set as a closed-loop drug delivery system based on ongoing neuronal activity detection, fluctuations in physiological recordings, or ongoing behavior. By making intravenous drug delivery fully programmable rather than mechanically constrained, WEARIT establishes a technological framework capable of evolving alongside future advances in behavioral neuroscience and systems neurobiology. Through future integration with an IoT-based wireless-network system, the Bluetooth-enabled architecture could also be extended to support remotely programmable and coordinated operation across larger numbers of devices, extending closed-loop control from individual animals to group-housed or multi-animal experimental paradigms^25,28,38^.

In summary, WEARIT establishes more than a wireless replacement for conventional tethered infusion systems. It provides a versatile technological platform for temporally precise intravenous pharmacology in freely behaving animals, enabling experiments that were previously difficult to perform. By combining wireless drug delivery with unrestricted behavior, complex environments, and state-of-the-art neuronal recording and manipulation approaches, WEARIT can enable new opportunities to investigate the neuronal mechanisms of drug action, reinforcement, circuit function, and therapeutic interventions across a broad range of disciplines extending from addiction neuroscience to systems neuroscience, neuropharmacology, and translational drug development.

## METHODS

### Fabrication of the WEARIT device

#### Drug reservoir

The drug reservoirs were fabricated from polyethylene glove material and silicone tubing (**Supplementary Fig. 3**). Individual finger sections were cut from the glove and heat-sealed on three sides using a heat sealer (K1, Reforest) at 85 ℃ for 5 s, repeated three times, to form a U-shaped pouch containing a total volume of 5 mL. The remaining open end was trimmed with scissors, and a silicone tube (outer diameter (OD): 3 mm, inner diameter (ID): 1 mm, length: 30 mm; mp-s, Bartels Mikrotechnik) was inserted into the pouch. The two sides flanking the tubing were heat-sealed at 85 ℃ for 5 s, repeated twice, and the remaining unsealed portion of the opening was sealed with marine adhesive (3M Marine Adhesive Sealant 5200, 3M).

### Design and construction of the wireless infusion module

The wireless infusion module (**Fig. 1c**) was developed to deliver programmable fluid volumes to freely behaving rats by integrating wireless control electronics, a piezoelectric micropump, a rechargeable battery and a drug reservoir into a miniaturized wearable platform. The wireless circuit consisted of a 3.5 V low-dropout voltage regulator (NCP161BFCS350T2G, onsemi), three n-channel MOSFETs (RUM002N02T2L, ROHM Semiconductor) configured as a switch array for flow-rate control, and a BLE SoC (EYSHSNZWZ, Taiyo Yuden) for wireless communication. The control circuit was constructed on a flexible printed circuit board (fPCB) to achieve a lightweight, compact form factor. The fPCB was designed using Altium Designer (Altium) and fabricated by a commercial vendor (PCBWay). The design enabled folding of the circuit within the enclosure and allowed the extended charging pad to be stowed inside the housing during operation while remaining externally accessible during battery charging. A piezoelectric micropump (mp6-liq, Bartels Mikrotechnik) was connected to the circuit by soldering its connector to the fPCB using solder paste (SMDLTLFP10T5, Chip Quik) and was driven by a dedicated driver module (mp6-OEM, Bartels Mikrotechnik) embedded in the control circuit. Multiple infusion rates (1.05 mL min^-1^, 1.26 mL min^-1^, and 1.55 mL min^-1^) were achieved through software-controlled switching of the MOSFET array. The system was powered by a rechargeable 120 mAh lithium-polymer battery (JA367, Coms), which was charged via a custom docking charger (**Supplementary Fig. 4**) without requiring enclosure disassembly. The drug reservoir was connected directly to the micropump inlet to eliminate the dead volume associated with external infusion lines.

### Design, 3D printing, and assembly of the modular wearable enclosure

The modular wearable enclosure consisted of three main components: a replaceable drug cartridge, a wireless module housing, and a quick-release dock. Three-dimensional CAD models were designed using Autodesk Inventor (Autodesk), and all components were fabricated by fused filament fabrication (FFF)-based 3D printing (Bambu Lab P1S, Bambu Lab) with polylactic acid (PLA) filament. An adjustable Velcro strap (width: 38 mm, length: 25 cm; VS3810, YMCRLUX) was threaded through the strap slot of the quick-release dock, allowing the fit to be tailored to animals with different body sizes. Cylindrical neodymium magnets (diameter: 5 mm; thickness: 1 mm; ND 5×1, Neez) were embedded in the drug cartridge and the wireless module housing to provide tool-free attachment and detachment. The wireless control circuit, rechargeable LiPo battery, micropump, and drug reservoir were assembled within the enclosure (**Supplementary Fig. 2**), which provided mechanical protection and enabled backpack-style mounting on the animal.

### Fluidic interface for intravenous access

The micropump outlet was connected to silicone tubing (OD: 3 mm, ID: 1 mm, length: 100 mm; mp-s, Bartels Mikrotechnik) coupled to a Luer lock connector (RSN041, BleedZone), which interfaced with a rat vascular access harness (VAHR1H/22, Instech). This configuration established a fluidic connection between the wearable device and an intravenous catheter. A silastic catheter (C30PU-RJV2329, Instech) was surgically implanted into the jugular vein (see surgical procedure details in this methods section) and connected to the harness (VAHR1H/22, Instech), providing an intravenous access route for drug infusion in freely behaving rats.

### Design of the wireless lever-triggered operant system

A custom wireless triggering system was developed to enable untethered operant behavioral experiments, providing real-time communication between a conventional operant chamber and the wearable infusion device. The system consisted of a central controller integrated with the operant chamber and a battery-powered peripheral controller embedded within the wearable platform.

The operant chamber was equipped with a lever connected to a TTL interface module (TTL 28 V DC Adapter, Med Associates), which converted lever-press events into transistor–transistor logic (TTL) signals routed to the digital input pins of a central controller (nRF52840 Development Kit, Nordic Semiconductor). Upon detection of a rising-edge TTL event, the central controller transmitted a predefined command packet to a designated peripheral device (WEARIT) via Bluetooth Low Energy (BLE). Operating as a BLE central device, the controller maintained simultaneous connections with multiple WEARITs, with each input channel assigned a unique command identifier to enable selective activation of individual devices.

The WEARIT platform featured a peripheral controller (EYSHSNZWZ, Taiyo Yuden) functioning as a BLE peripheral. Upon receiving a command packet matching its assigned identifier, the peripheral controller generated a digital output signal to activate the drug-delivery circuitry. To prevent unintended repeated dosing, any lever press occurring during an active infusion or within 20 s after infusion completion was ignored. Output signal duration and the activation sequence were defined by embedded firmware, ensuring reproducible operation after each lever press.

### Validation of infusion volume

Infusion volume was validated gravimetrically. Delivered fluid was collected in a sealed 10 mL vial fitted with a 20-gauge catheter inserted through the cap; the insertion point was sealed to prevent evaporation and leakage. The vial was weighed before and after infusion using a microbalance (XSR105DUV, Mettler Toledo), and the mass difference was converted to volume assuming the density of water. The measured volume was compared with the programmed target volume to assess delivery accuracy.

### Animals

All procedures were approved by University of California, Los Angeles Animal Care and Use Committee in accordance with the National Institutes of Health Guidelines for the Care and Use of Laboratory Animals. Adult male and female Long Evans Wild Type (250–350 g) were used for this study. All animals for behavioral experiments were 8 to 10 weeks old at the beginning of the experiments. Rats were group housed with two to three animals per cage on a 12/12 hours dark/light cycle (lights on at 7:00 AM) and acclimated to the animal facility holding rooms for at least 7 days before any manipulation. All experiments were performed during the light cycle. Rats received food and water *ad libitum* until 2 days prior to starting the food self-administration behavioral studies, when food restriction (16 g of rat chow per day) started and continued until the end of the experiments.

### Surgeries

All surgeries were performed under isoflurane (2.5/3 MAC) anesthesia using appropriate sterile aseptic techniques, as published previously^36,47–49^. After confirming sedation, small incisions were made both on the dorsal surface of the neck and on the ventral surface of the neck. On the latter, the underlying tissue was carefully dissected to expose and isolate the jugular vein. A sterile indwelling catheter (C30PU-RJV2329, Instech) was inserted into the jugular vein and sutured in place using non-absorbable silk suture (MedVet International). The catheter was then tunneled subcutaneously using a buck ear curette (Sklar, Blunt tip #3) and exited the body through the first small hole in neck on the dorsal side. The ventral incision was closed using non-absorbable silk suture (MedVet International). The exposed catheter was connected to a One Channel Vascular Access Harness for rats (VAHR1H/22, Instech) containing a 22 ga port for drug administration. Animals were given Carprofen (2 mg kg^-1^ s.c.), Baytril (8 mg kg^-1^ s.c.), and bupivacaine (5 mg kg^-1^, site of incision). In addition, rats were given Carprofen tablets (Bio Serv, MD150-2) for 2 days after surgery to assist in wound healing and analgesia. Due to the nature of the vascular harness, rats were singly housed following surgery and were allowed to recover for 1 week prior to random assignment to an experimental group. Catheter patency was maintained with daily flushing of 0.3 mL sterile saline containing gentamicin (1.33 mg mL^-1^, i.v.). Rats with a loss of catheter patency were excluded from the study.

### Respiratory depression experiment

To monitor oximetry on freely moving rats receiving fentanyl (50 µg kg^-1^) from WEARIT device, we used a non-invasive pulse oximeter collar (Starr LifeSciences), as described previously^36,47,50^. Oxygen saturation was collected (MouseOx v2.0, Starr LifeSciences) before and after an experimenter-induced bolus infusion of intravenous fentanyl, or saline as a control, as a proxy for respiratory depression^36,47^.

### Operant conditioning

#### Sucrose

Sucrose self-administration was conducted using operant-conditioning chambers (Med Associates, MED – PC 5 software) equipped with two retractable levers positioned on the right-hand wall and a food magazine connected to a food pellet dispenser. Two cue lights were positioned above the levers, and one house light was positioned on the top left-hand wall. During self-administration sessions both levers (active and inactive) were extended out with white cue light turned on only above active lever. Presses on the active lever resulted in delivery of a 45 mg chocolate flavored sucrose pellet (F0025, BioServ) and a 20 s time-out period during which active and inactive lever were retracted, and cue light above active lever was turned off. Presses on the inactive lever had no consequences and were used to determine accurate learning of the operant task.

For learning procedures, animals were food restricted and were fed daily with a total of 10% of their weight in food pellets. Animals overall weight was monitored during the whole experiment to ensure that rats maintained a stable weight. Rats were habituated to carry the WEARIT device on their back for at least 15 minutes daily for a minimum of 5 days. During that period, rats were also exposed to a daily 1 hour Fixed Ratio (FR) 1 schedule of reinforcement for sucrose self-administration session, which was ran before daily habituation to the device. Appropriate learning was defined as 3 consecutive daily sessions where active levers presses represented 75% of total presses. All animals achieved this in 5 sessions. A day after learning completion rats were equipped with either a harness (control) or WEARIT device and placed in operant boxes for a 1 hour sucrose self-administration session. The following day, conditions (harness only vs WEARIT) were counterbalanced for each animal, allowing for intra-individual control of performances. Active and Inactive lever presses, as well as number of pellets obtained during these sessions were monitored.

#### Fentanyl

A week after catheter implantation (see procedure details above), all rats were habituated to carrying the WEARIT device for at least 15 minutes daily for five days. After habituation, rats were either tethered to receive fentanyl from a classic pump ^1,47–49,51^ or equipped with the WEARIT device, loaded with 5 mL of fentanyl. Fentanyl concentration was adjusted to deliver constant doses per injection for each individual animal (5, 2.5, and 1 µg kg^-1^). Briefly, rats were placed in the operant chambers for an hour and active and inactive lever presses, and number of fentanyl infusions were monitored. For the first five sessions, rats received a dose of 5 µg kg^-1^ of fentanyl, as described before ^52^. After five days, fentanyl concentrations were decreased to 2.5 µg kg^-1^ for five additional days, before being lowered to 1.25 µg kg^-1^ for five days. This strategy allowed us to assess a dose response for fentanyl consumption ^51^ and compare consumption between the classic tethered and WEARIT intravenous self-administration approaches. Catheters were flushed daily with 0.3 mL sterile saline containing gentamicin (1.33 mg mL^-1^, i.v.) before and after self-administration sessions to ensure catheter patency.

### Open field test for locomotor activity

Open field test was performed in a black enclosure (80 × 80 cm^2^) covered with fresh bedding. Rats’ locomotion was monitored and analyzed using a videotracking software (AnyMaze, Stoelting). Rats were exposed to daily 20-minutes habituation sessions for five days during which they were equipped with the WEARIT device. A day after the last habituation session, rats were equipped with either harness alone, or the WEARIT device and placed in the open field for 20 minutes. Horizontal activity was monitored.

In a separate experiment, animals were habituated to the WEARIT device for 3 days, as described above. The following day, animals were injected with Amphetamine (2 mg kg^-1^) to assess whether WEARIT would impact stimulant-induced hyperactivity.

### Quantification and Statistical Analysis

All experiments were performed in at least two separate cohorts of rats and all treatment groups in each cohort were performed at least twice to avoid any nonspecific day/condition effect. Animals were randomly assigned to WEARIT or Control - or saline vs Fentanyl for oximetry experiments - groups before testing. Statistical significance was taken as * *P* < 0.05, ** *P* < 0.01, *** *P* < 0.001, and **** *P* < 0.0001, as determined by a one-way ANOVA or a two-way repeated-measures ANOVA followed by a Sidak post-hoc tests, two-tailed unpaired or paired t-test, or Mann-Whitney for unpaired values as appropriate. All data samples were tested for normality of distribution using Shapiro-Wilk test before being assigned to ANOVAs, two-tailed t-test, two-tailed Mann-Whitney for unpaired values, or two-tailed Wilcoxon matched-pairs signed rank analysis. All data are expressed as mean ± s.e.m. Sample size (n number) always refers to the value obtained from an individual animal when referring to behavioral experiments. Statistical analyses were performed in GraphPad Prism 9.0 and SPSS.

## ACKNOWLEDGEMENTS

We thank all members of the Jeong, Massaly, and McCall labs for their insights, comments, and contributions to the development of this project, including early conversations with Marie Walicki.

## AUTHOR INFORMATION

## Contributions

J.G.M., J-W.J., and N.M. conceived, designed the study, and supervised all the work. E.Y.J., C.T., A.R., and N.M. performed the experiments, collected, processed and analyzed data. S.C., J.L., T.A., K.L., M.N., T.L., and E.S. contributed to the data collection and acquisition. E.Y.J., J-W.J., and N.M. created the figures and wrote the paper. E.Y.J., K.E.P., C.M.C., J.G.M., J-W.J, and N.M. edited the paper. All authors reviewed the paper and approved the decision to submit for publication.

## ETHICS DECLARATIONS

