## Supplemental Figures 1-8 & Supplemental Table 1 for "A Wireless Wearable Platform for Intravenous Drug Self-Administration in Freely Behaving Rats"

### SUPPLEMENTARY FIGURES & LEGENDS

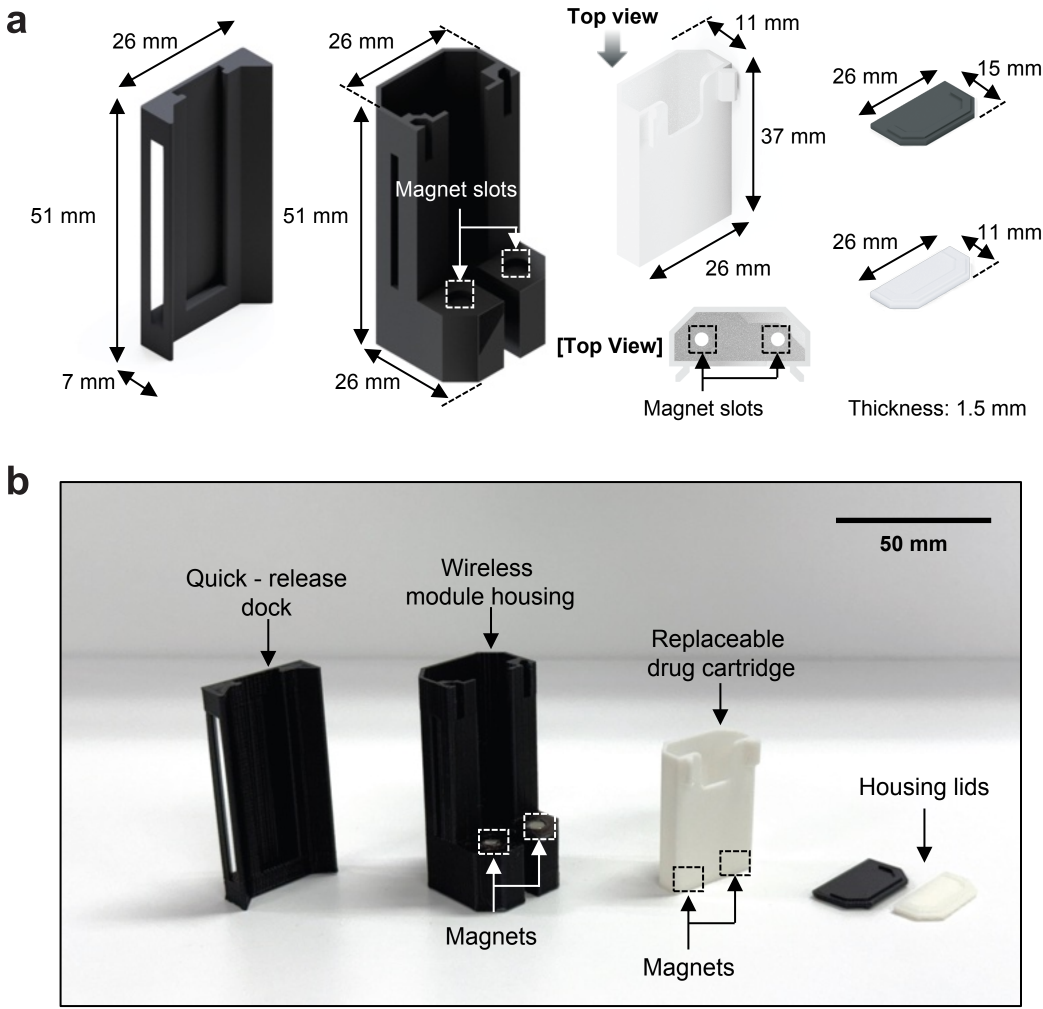
a
26 mm
51 mm
7 mm
26 mm
51 mm
Magnet slots
26 mm
Top view
11 mm
37 mm
26 mm
[Top View]
Magnet slots
26 mm
15 mm
26 mm
11 mm
Thickness: 1.5 mm
b
50 mm
Quick - release dock
Wireless module housing
Replaceable drug cartridge
Housing lids
Magnets
Magnets

**Supplementary Fig. 1 | Design and fabrication of the 3D-printed enclosure. a**, Dimensioned renderings of the modular enclosure components, including the quick-release dock, wireless module housing, replaceable drug cartridge, and housing lids for the wireless module housing and drug cartridge, with corresponding dimensions. Magnet slots in the wireless module housing and replaceable drug cartridge accommodate embedded neodymium magnets, enabling modular magnetic attachment. **b**, Photograph of the fabricated enclosure components. Neodymium magnets embedded within the wireless modular enclosure components enable rapid assembly/disassembly and cartridge replacement.

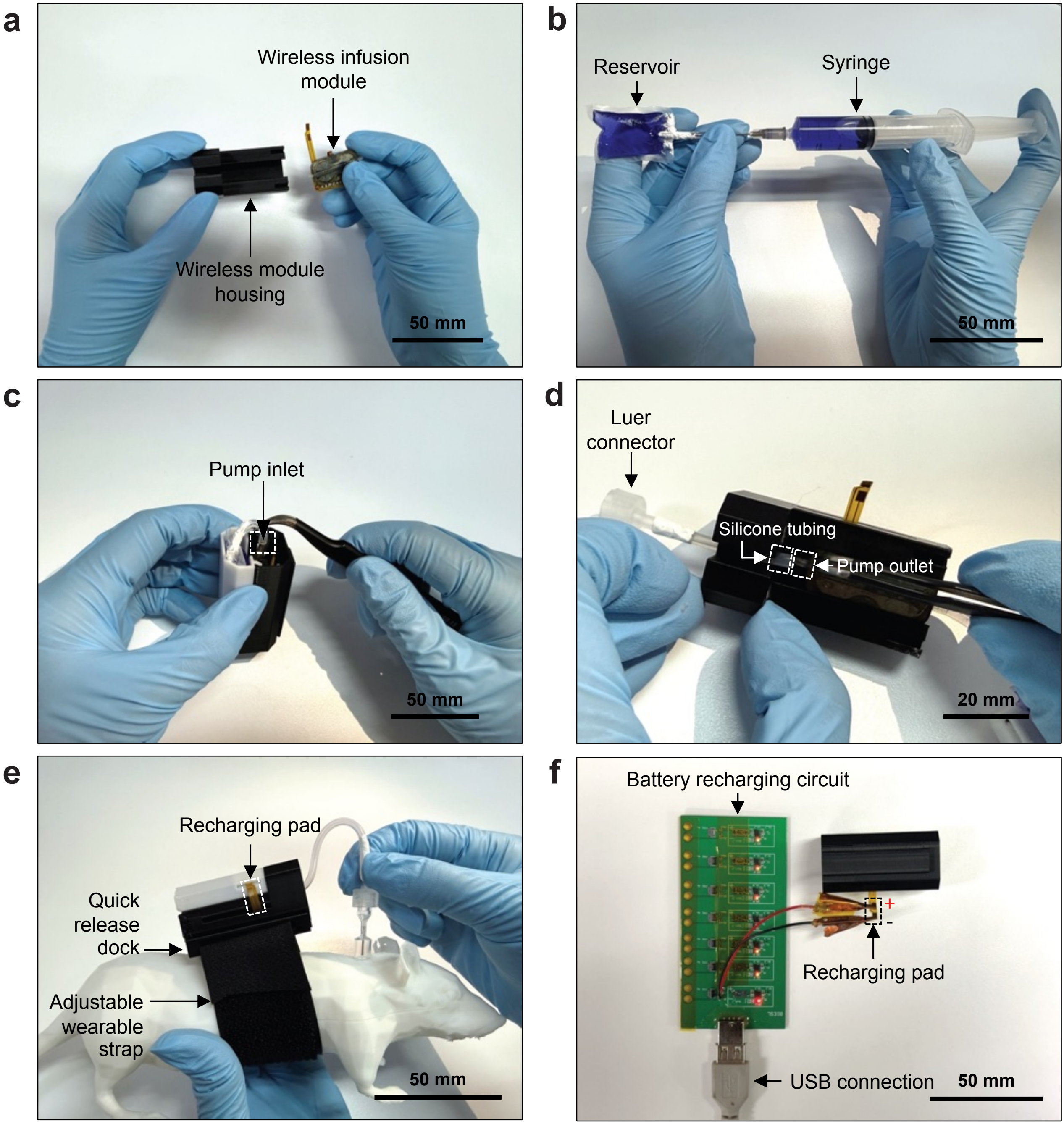
a
Wireless infusion module
Wireless module housing
50 mm
b
Reservoir
Syringe
50 mm
c
Pump inlet
50 mm
d
Luer connector
Silicone tubing
Pump outlet
20 mm
e
Recharging pad
Quick release dock
Adjustable wearable strap
50 mm
f
Battery recharging circuit
+
-
Recharging pad
USB connection
50 mm

**Supplementary Fig. 2 | Assembly procedure of the WEARIT device. a**, The assembled wireless infusion module was inserted into the wireless module housing. **b**, The replaceable drug reservoir was filled using a syringe prior to assembly. **c**, The filled drug reservoir was placed into the reservoir protective cartridge, and the reservoir outlet tubing was connected to the micropump inlet. **d**, The micropump outlet was connected to silicone tubing and a Luer-lock connector for connection with the rodent harness. **e**, The assembled wireless module was mounted onto the quick-release dock. The fluidic interface is coupled to the vascular access harness to establish intravenous access. The recharging pad is protected within the 3D-printed enclosure during freely moving behavior. **f**, The recharging pad enables connection to a custom USB-powered battery charging circuit without disassembling the modular enclosure.

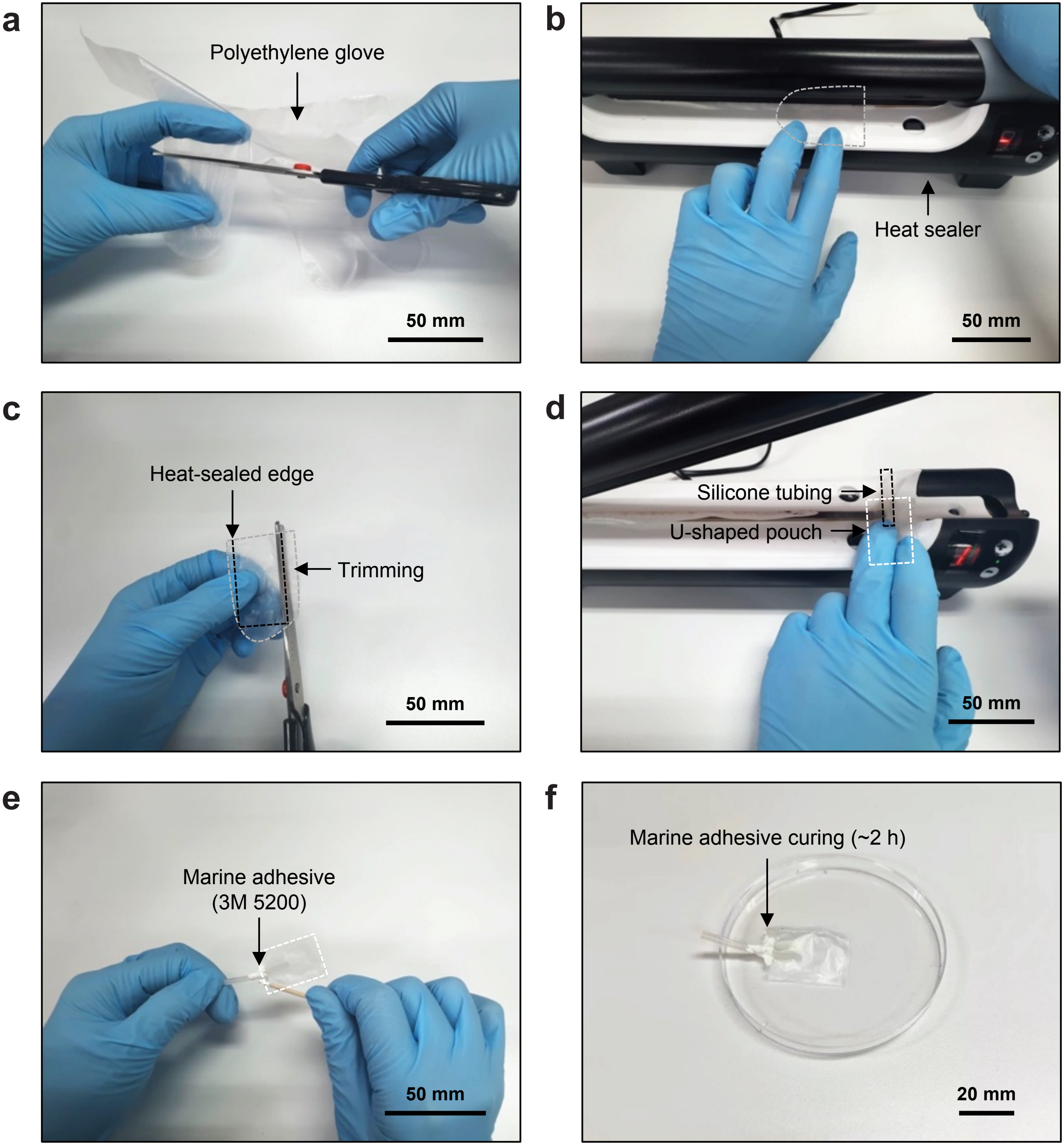
a
Polyethylene glove
50 mm
b
Heat sealer
50 mm
c
Heat-sealed edge
Trimming
50 mm
d
Silicone tubing
U-shaped pouch
50 mm
e
Marine adhesive
(3M 5200)
50 mm
f
Marine adhesive curing (~2 h)
20 mm

**Supplementary Fig. 3 | Fabrication process of the replaceable drug reservoir. a**, Individual sections were cut from a polyethylene glove to form the reservoir body. **b**, Three sides of the polyethylene pouch were heat-sealed using a heat sealer to create a closed U-shaped reservoir structure. **c**, The heat-sealed edges were trimmed to remove excess material. **d**, A silicone tube was inserted into the unsealed fourth side of the reservoir, and the two sides adjacent to the tubing were then heat-sealed. **e**, The remaining unsealed section was sealed with marine adhesive (3M Marine Adhesive Sealant 5200) to prevent leakage. **f**, The adhesive was cured for ~2 h to complete the drug reservoir.

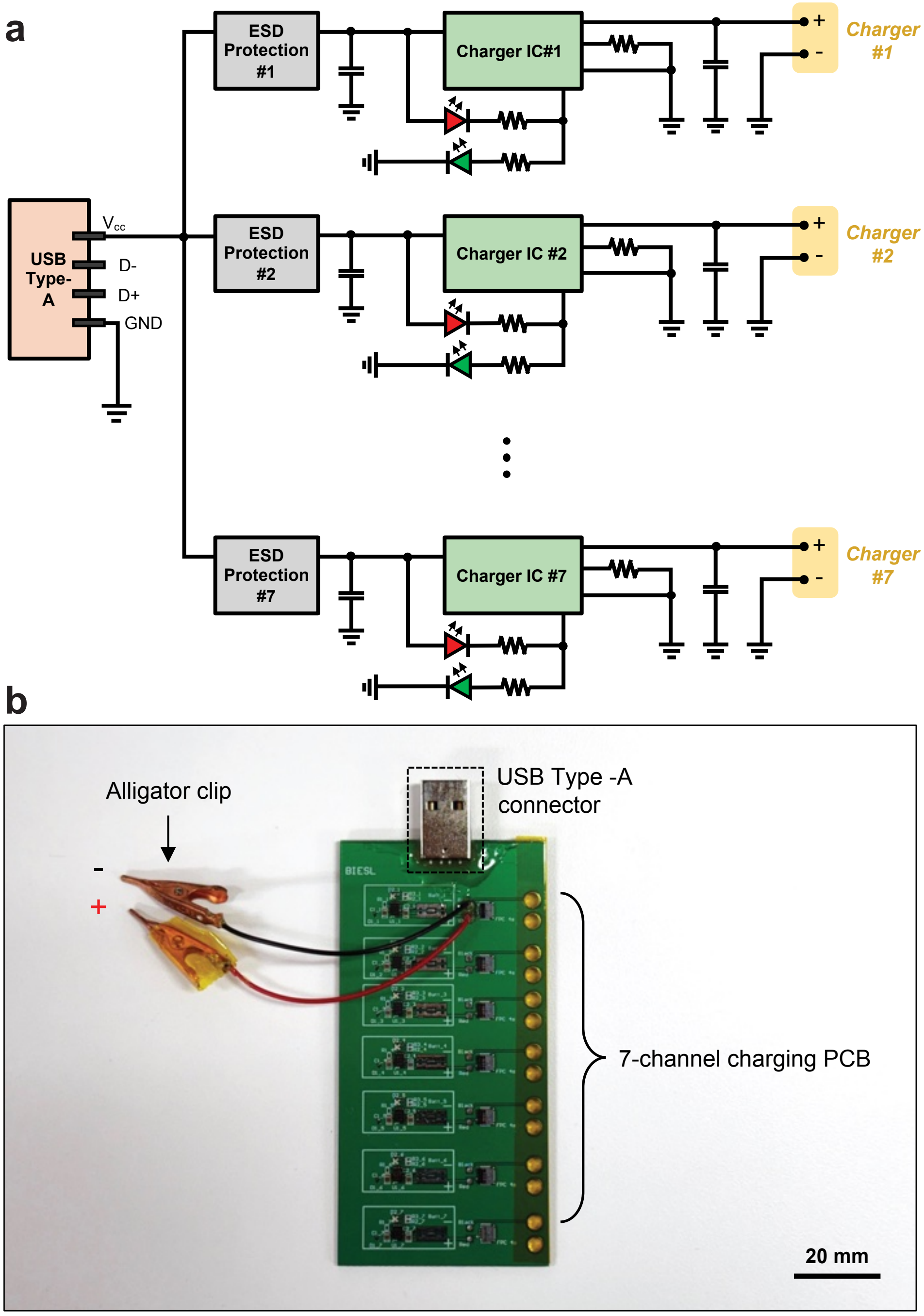
a
USB
Type-
A
Vcc
D-
D+
GND
ESD
Protection
#1
Charger IC#1
Charger
#1
ESD
Protection
#2
Charger IC #2
Charger
#2
ESD
Protection
#7
Charger IC #7
Charger
#7
b
Alligator clip
-
+
USB Type -A
connector
BIESL
7-channel charging PCB
20 mm

**Supplementary Fig. 4 | Custom multi-channel battery charger for WEARIT devices.** **a**, Circuit schematic of the custom charging PCB. A USB Type-A connector supplies power to seven independent charging channels. Each channel consists of an ESD protection circuit, battery charger IC, status-indicator LEDs, and passive filtering components, enabling simultaneous charging of up to seven WEARIT devices. **b**, Picture of the fabricated 7-channel charging PCB. An alligator clip is shown connected to one representative output channel; each of the seven channels can be used independently to charge additional WEARIT devices.

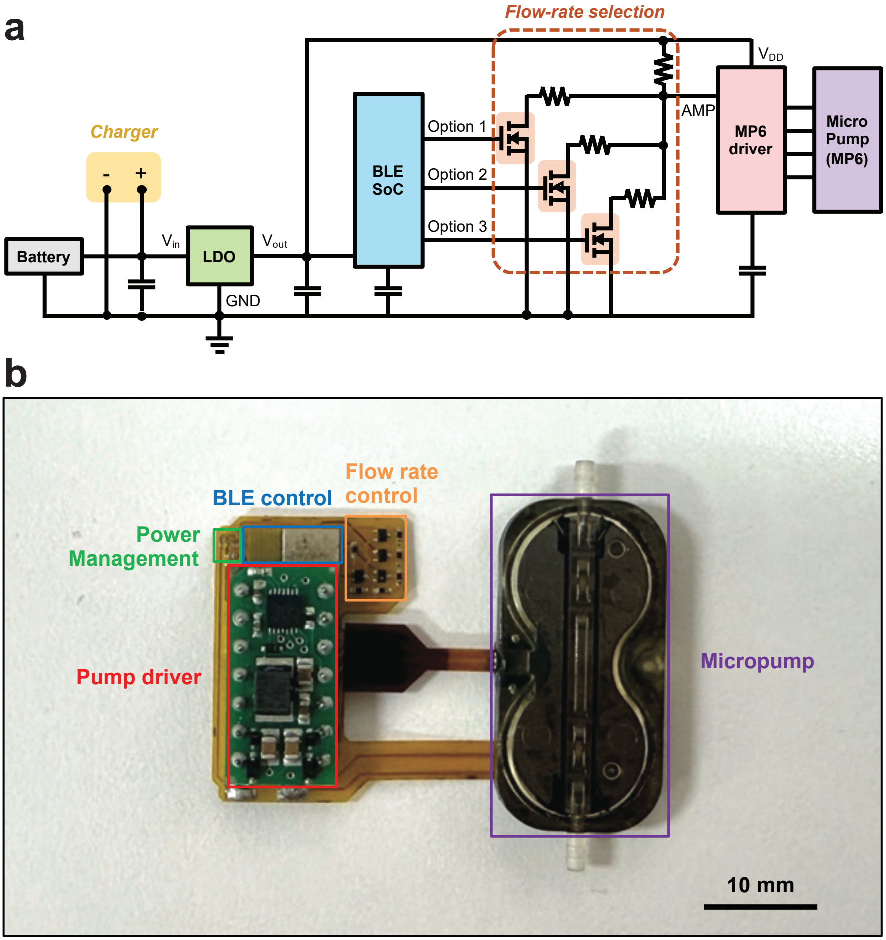
a
Flow-rate selection
Charger
-
+
Battery
Vin
LDO
Vout
GND
BLE SoC
Option 1
Option 2
Option 3
VDD
AMP
MP6 driver
Micro Pump (MP6)
b
Power Management
BLE control
Flow rate control
Pump driver
Micropump
10 mm

**Supplementary Fig. 5 | Circuit architecture and implementation of the wireless infusion module. a**, Circuit schematic of the wireless infusion module. Power from the rechargeable battery is regulated by a low-dropout regulator (LDO) and supplied to the BLE system-on-chip (SoC) and micropump driver. Software-selectable flow-rate control is achieved through flow-rate selection circuitry (i.e., MOSFET-based switch array) that adjusts the pump-driver input, enabling multiple infusion-rate settings. **b**, Picture of the assembled wireless infusion module, with functional blocks – power management, BLE control, flow-rate control, pump driver, and micropump – corresponding to the circuit elements in **a**.

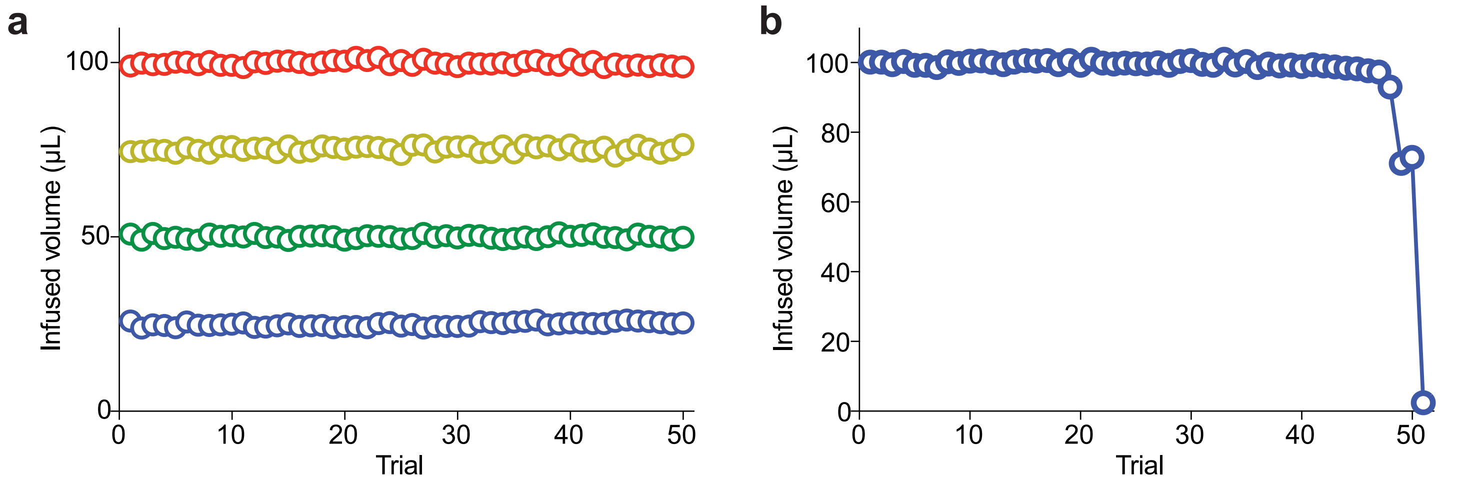
a
b
Infused volume (μL)
Trial
100
50
0
10
20
30
40
80
60

**Supplementary Fig. 6 | Reproducibility of programmed infusion volumes and reservoir depletion characterization. a**, Repeated infusion tests performed using an external fluid reservoir connected directly to the micropump inlet. Programmed infusion volumes of 25, 50, 75, and 100 μL were delivered over 50 consecutive trials, demonstrating stable and reproducible volume delivery across all tested settings. **b**, Reservoir depletion test using the 5 mL wearable drug reservoir, with the infusion volume programmed to 100 μL per actuation and the delivered volume monitored across consecutive infusion cycles. Delivery accuracy remained within 2.67% of the target volume through the 48th infusion, after which accuracy declined as the reservoir approached depletion.

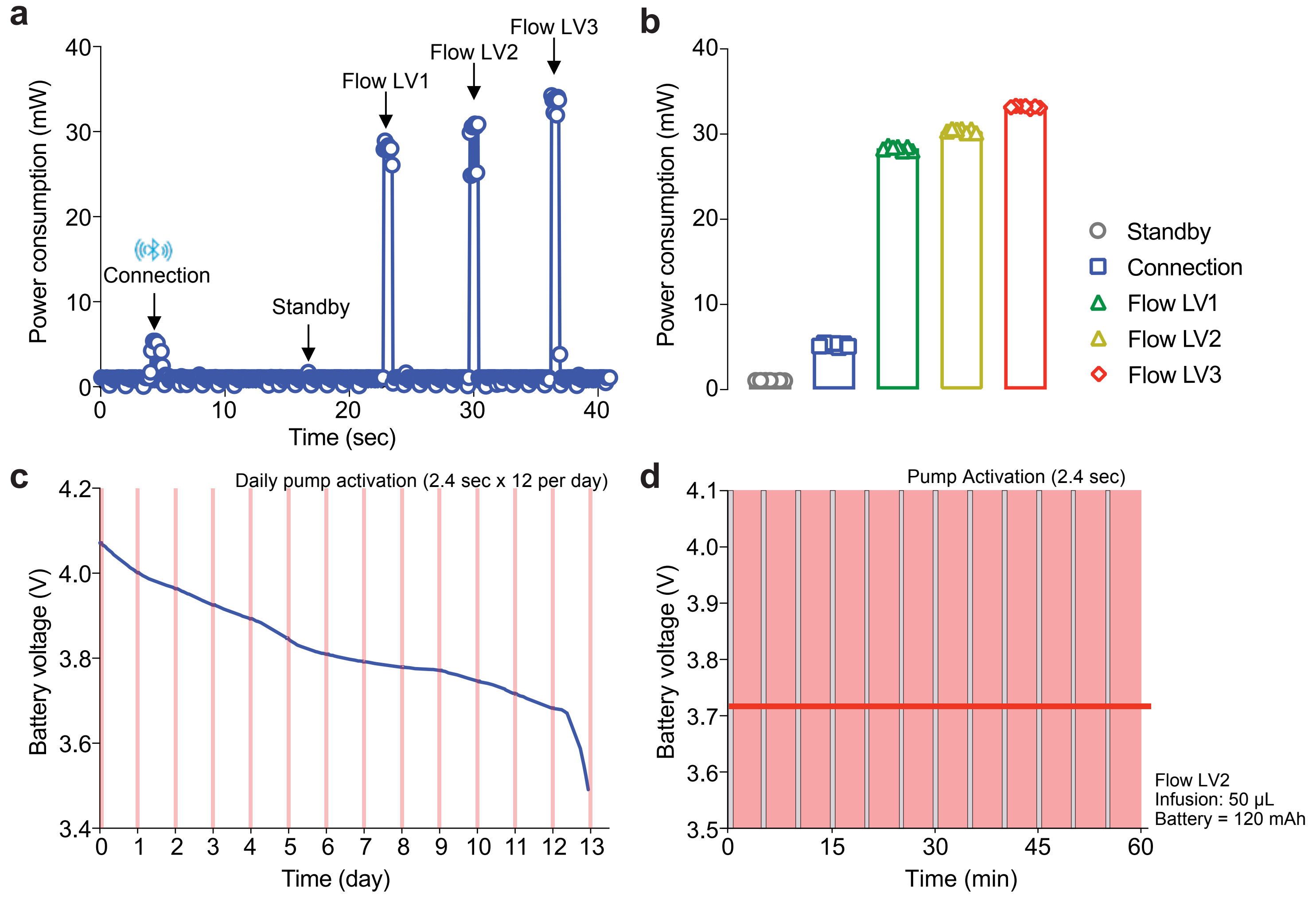
a
Power consumption (mW)
Time (sec)
Connection
Standby
Flow LV1
Flow LV2
Flow LV3
b
Power consumption (mW)
Standby
Connection
Flow LV1
Flow LV2
Flow LV3
c
Daily pump activation (2.4 sec x 12 per day)
Battery voltage (V)
Time (day)
d
Pump Activation (2.4 sec)
Battery voltage (V)
Time (min)
Flow LV2
Infusion: 50 μL
Battery = 120 mAh

**Supplementary Fig. 7 | Power consumption and battery performance of WEARIT. a**, Representative power-consumption profile of the wireless infusion module during standby operation, Bluetooth Low Energy (BLE) connection, and micropump actuation at three flow-rate settings (Flow LV1–LV3). Power consumption increased transiently during wireless connection and pump operation, with higher flow-rate settings resulting in greater power demand. **b**, Average power consumption measured during standby operation, BLE connection, and pump operation at the three programmable flow-rate settings. Data are represented as mean ± s.d. ($n$ = 7 measurements). **c**, Battery lifetime characterization of the WEARIT device powered by a 120-mAh lithium-polymer battery. Battery voltage was monitored during repeated daily pump activation sessions (one hour per day) consisting of twelve 50 μL infusions delivered at Flow LV2 at 5-min intervals. The device remained operational for up to 13 days before battery voltage declined to 3.5 V, the device's operational threshold. **d**, Magnified view of the battery-voltage profile during a single one-hour pump-activation session, demonstrating minimal transient voltage fluctuations during repeated pump actuation.

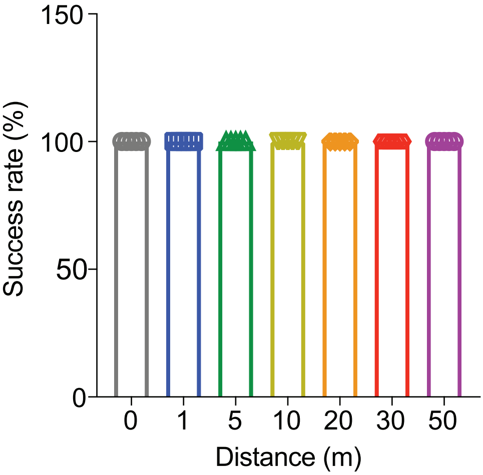

150
100
50
0
Success rate (%)
0
1
5
10
20
30
50
Distance (m)

**Supplementary Fig. 8 | Wireless communication success rate as a function of distance.** Success rate of wireless triggering from the central controller to the WEARIT device measured at separation distances of 0, 1, 5, 10, 20, 30, and 50 m under line-of-sight conditions. Reliable communication was maintained across all tested distances, with a 100% success rate under each condition (*n* = 5 trigger events per distance).

| Category | Component | Supplier | Model number | Unit cost (USD) | Subtotal (USD) | Total (USD) |
|---|---|---|---|---|---|---|
| Wireless infusion module | Piezoelectric micropump | Bartels Mikrotechnik | mp6-liq | 76.70 | 205.69 | 421.64 |
| | Pump driver | Bartels Mikrotechnik | mp-Driver | 94.90 | | |
| | Silicone tubing | Bartels Mikrotechnik | mp-s | 10.40 | | |
| | Flexible control PCB | PCBWay | Custom flexible PCB | 1.29 ($258 for 200 pcs) | | |
| | Bluetooth SoC | Taiyo Yuden | EYSHSNZWZ | 20.00 | | |
| | LiPo battery | Coms | JA367 | 2.40 | | |
| Operant-triggered wireless control system | BLE central controller | Nordic Semiconductor | nRF52840 Development Kit | 48.95 | 215.95 | |
| | TTL interface module | Med Associates | TTL 28 V DC Adapter | 167 | | |

**Supplementary Table 1 | Components and costs of the WEARIT system.** Major components used in the system and their approximate unit costs.
